# Cell cycle-balanced expression of pluripotency regulators via cyclin-dependent kinase 1

**DOI:** 10.1101/764639

**Authors:** Sergo Kasvandik, Reelika Schilf, Merilin Saarma, Mart Loog, Kersti Jääger

## Abstract

Embryonic stem cells (ESCs) have a unique ability to remain pluripotent while undergoing rapid rounds of cell division required for self-renewal. However, it is not known how cell cycle and pluripotency regulatory networks co-operate in ESCs. Here, we used stable isotope labeling with amino acids in cell culture (SILAC) combined with mass spectrometry to determine pluripotency proteome dynamics during cell cycle in mouse ESCs. We found the S/G2M-fluctuating pluripotency transcription factors (ESRRB, REST), chromatin regulators (JARID2, TRIM24) and proteins with E3 ligase activity (NEDD4L, PIAS2) to peak in S phase. This expression balance was disrupted upon inhibition of cyclin-dependent kinase 1 (CDK1) activity resulting in the shift of the expression peak from S to G2M. Our results demonstrate that mouse ESCs require CDK1 activity to maintain high S to G2M ratio of pluripotency regulators revealing critical role of cell cycle dynamics in balancing ESC identity.

## INTRODUCTION

Embryonic stem cells (ESCs) are the best studied pluripotent stem cell system that encompasses the abilities of self-renewal and differentiation. How this ability is manifested during cell division cycle is still unclear. Mouse ESCs (mESCs) grown in serum/LIF culture conditions proliferate rapidly and spend most of their time in S phase due to untypically short G1 phase (Stead et al., 2002). The pluripotency transcription factors (TFs) including OCT4, NANOG and ESRRB (van den Berg et al., 2008; Chambers et al., 2003; Nichols et al., 1998; Niwa et al., 2000) govern the intrinsic regulatory network that via interaction with extrinsic cues determine cell fate. However, our knowledge about ESC decision-making using bulk populations best reflects TF expression and interactions present in the more dominant S phase.

Connection of pluripotency TF network with cell cycle occurs via cyclin-dependent kinase 1 (CDK1) that along with Cyclin B are key players in cell cycle regulation and particularly mitosis in eukaryotic cells. CDK1 is indispensible for the early development of the embryo, and is required for the self-renewal of mESCs and human ESCs (Neganova et al., 2014; Zhang et al., 2011). CDK1 inhibition has been shown to trigger abnormal transcription of OCT4 target genes in mitosis and lead to loss of pluripotency in mESCs (Kim et al., 2018).

Some studies have reported relatively uniform expression of pluripotency TFs during cell cycle both at mRNA and protein level (Shin et al., 2016; Singh et al., 2013), while others have revealed fluctuations (Gonzales et al., 2015; Van Der Laan et al., 2014; Tsubouchi et al., 2013). Given the interconnection between ESC fate choices and cell cycle (Gonzales et al., 2015; Jääger et al., 2019; Van Oudenhove et al., 2016; Pauklin and Vallier, 2013; Petruk et al., 2017), we aimed to map mESC proteome dynamics during cell cycle on a global scale.

We used stable isotope labeling with amino acids in cell culture (SILAC) combined with mass spectrometry (MS)-based identification of proteins in G1-, S- and G2M-enriched mESCs both in untreated and Cdk1-inhibitor treated cells to reveal cell cycle coordinated dynamics of pluripotency factor expression. Our experimental design enabled quantification of ca 2500 proteins across cell cycle phases and identify major reorganisation of pluripotency network expression at S-to-G2M transition upon loss of CDK1 activity.

## EXPERIMENTAL PROCEDURES

### mESC culture and stable isotope labeling with amino acids in cell culture (SILAC)

Cell cycle reporter N7-mESCs that express Cyclin B1:GFP fusion protein from an endogeneous locus (Jääger et al., 2019) were cultured on 0.1% gelatin-coated tissue culture plastic in GMEM/β-mercaptoethanol/10%FCS/LIF as described before (Smith, 1991). For stable isotope labeling with amino acids in cell culture (SILAC), N7-mESCs were cultured for two weeks in DMEM/β-mercaptoethanol/LIF w/o Arg and Lys (Gibco), supplemented with 12.5% KnockOut™ Serum Replacement (Gibco), 63 mg/L light-Arg and 147 mg/L light-Lys or 147 mg/L heavy (13C615N2)-Lys (Lys8) (Cambridge Isotope Laboratories). N7-mESCs cultured in SILAC-media were viable and proliferated normally as evidenced under microscope at passages 3 and 5 (**Figure S1A**).

### Cell cycle sorting of labelled cells

Light-Lys (L) or heavy-Lys (H) labelled N7-mESCs were sorted into different cell cycle phase fractions using flow cytometry as in (Jääger et al., 2019). Briefly, N7-mESCs were stained with 50 uL Hoechst 33342 (50 mg/mL stock) in 10 mL SILAC medium 45 min at 37°C prior to FACS-sorting using the indicated gates (**Figure S1B**). Sorted cells (on average 500 000 cells per fraction) were immediately pelleted and stored at −80°C until protein sample preparation. CDK1 activity was inhibited in heavy-Lys (H) labelled N7-mESCs in SILAC/Hoechst medium prior to FACS sorting (2h in total) using 10 uM RO3306 (CDK1i, Axon 1530).

### Sample preparation for mass-spectrometry (MS) analysis

Altogether, we combined seven sorted N7-mESCs samples in two configurations. For proteome profiling in G1, S and G2M, light-Lys G1 or G2M were each mixed with heavy-Lys S phase sample 1:1 by cell number, resulting in two dual-SILAC samples G1-S and G2M-S (Setup 1, **Figure 1A**); for proteome profiling in G1, S and G2M each treated with CDK1i, we mixed heavy-Lys G1 or S or G2M each with light-Lys supermix (SM, containing equal amount of G1-S-G2M) 1:1 by cell number, which resulted in three dual-SILAC samples G1-CDK1i-SM, S-CDK1i-SM, G2M-CDK1i-SM (Setup 2, **Figure 1A**). Detailed description of the proteomic methods can be found in **Supplemental Information**. Briefly, cell pellets were lysed with heat and sonication in an SDS lysis buffer. Extracted proteins were methanol-chloroform precipitated and digested in denaturing conditions with Lys-C and trypsin. Peptides were desalted and injected to a Q Exactive Plus (Thermo Fisher Scientific) nano-LC MS/MS system operated in a data-dependent acquisition mode. MS raw data were identified and quantified with the MaxQuant software. Profiling and enrichment analysis of proteomics data is described in **Supplemental Information**.

**Figure 1.**
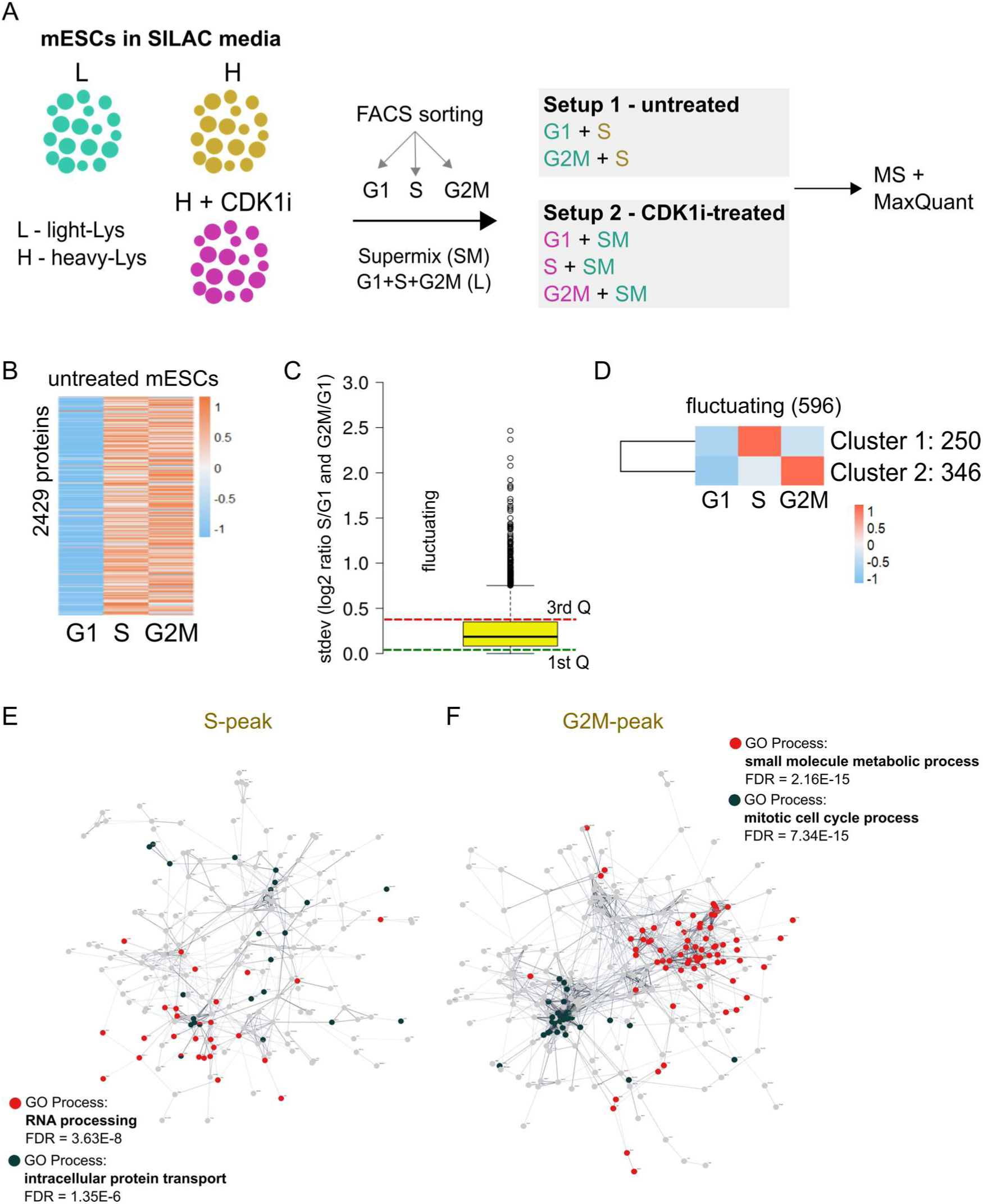
mESC proteome dynamics during cell cycle. **A**. A schematics showing the strategy of <u>s</u>table isotope labeling with <u>a</u>mino acids in <u>c</u>ell culture (SILAC) used in combination with mass spectrometry (MS) to determine proteome dynamics during cell cycle in mouse embryonic stem cells (mESCs). Cell cycle enrichment was performed using flow cytometry sorting (**Figure S1B**). Two MS runs were performed using the indicated sample setups 1 (untreated) and 2 (CDK1i-treated). L = light lysine, H = heavy lysine, CDK1i – Cyclin-dependent kinase 1 inhibitor **B**. An unclustered heatmap showing the scaled abundances of 2429 identified proteins in mESCs across cell cycle phases. **C**. Boxplot diagram showing distribution of differences (stdev-s) between S and G2M. Proteins above the red line (upper 3rd Q) were defined as ‘fluctuating’ and proteins below the green line (lower 1st Q) were defined as ‘constant’. Proteins between the colored lines were defined as ‘moderately changing’. **D**. Clustered abundance heatmap showing the distribution of S/G2M fluctuating proteins in C into two major clusters. Clusters were named by expression profile: S-peak (Cluster 1), G2M-peak (Cluster 2). **E and F**. Protein interaction networks of either S- or G2M-peaking protein sets derived from Cytoscape/stringApp analyses using edge-weighted network layouts. Edges represent protein-protein associations based on both experimental and indirect evidence. Unassociated proteins were excluded (S-peak: 42 proteins; G2M-peak: 47 proteins). Note sparsely interconnected network of S-peaking proteins in contrast to dense clustering in G2M-peaking protein set. Highly enriched GO Process terms were derived via STRING enrichment analysis. See also **Figure S**1 and **Supplemental Table 1**.

## RESULTS

### mESC proteome dynamics during cell cycle

To measure relative protein abundances in mESCs during cell cycle, we enriched isotope-labelled cell cycle reporter N7-mESCs (Jääger et al., 2019) for G1, S and G2M phases using flow cytometry (**Figure S1B**), and prepared two dual-SILAC samples where S phase standard was combined with either G1 or G2M (**Figure 1A**). This strategy enabled G1-S-G2M normalisation via S phase and direct inter-sample comparison, resulting in relative quantification of 2429 proteins during mESC cell cycle (**see Experimental Procedures, Supplemental Table 1**).

First, by global profiling of protein expression during cell cycle, we determined profound increase in overall protein abundances both in S and G2M phases compared with G1 (**Figure 1B**). This pattern might reflect increase in cell size towards S/G2M. However, as the active maintenance of pluripotency occurs in S and G2M phases (Gonzales et al., 2015), we were particularly interested in determining dynamic changes at this transition. To enable direct comparison between S and G2M, we used G1 as a reference and computed log2 ratios for S/G1 and G2M/G1. Analysis of the mitotic signature proteins obtained from Cyclebase (Santos et al., 2015) confirmed gradually increasing expression towards G2M (>log2-fold increase, G2M versus G1)(**Figure S1C**), in accordance with known cell cycle dynamics.

To identify groups of proteins with characteristic S/G2M dynamics, we calculated standard deviations (stdev-s) between log2 ratios of S/G1 and G2M/G1 and used box plot diagram of stdev-s to classify proteins into fluctuating (upper 3rd quartile, Q), constant (lower 1st Q) and moderately changing (the middle 50%) (**Figure 1C, Supplemental Table 1**), a strategy previously used for comparing cell cycle fractionated proteomes (Carpy et al., 2014). We focussed our analysis on the fluctuating proteome (596 proteins) and aimed to determine the direction of the change. K-means correlation clustering into two major clusters revealed 250 proteins to peak in S (Cluster 1) and 346 proteins to peak in G2M (Cluster 2) (**Figure 1D, Supplemental Table 1**). This analysis shows that globally, slightly larger proportion of S/G2M fluctuating proteins (58%) peak in G2M.

Gene ontology (GO) analysis using DAVID (Huang et al., 2009) revealed that the S phase-peaking protein set (Cluster 1) was enriched for biological processes related to transcription and protein transport (**Figure S1D**), whereas processes overrepresented in G2M-peaking set (Cluster 2) included cell divison and oxidation-reduction process, in line with cell cycle features. Protein interaction network analysis using STRING (Snel et al., 2000; Szklarczyk et al., 2019) identified sparsely connected topology amongst S-peaking proteins (**Figure 1E**), while proteins peaking in G2M formed dense association clusters of ‘mitotic cell cycle process’ and ‘small molecule metabolic process’ (**Figure 1F, Supplemental Table 1**), in accordance with DAVID analysis (**Figure S1D**). Overall, these results show that while major increase in global proteome abundance occurs at G1-to-S transition, the expression levels change both from low to high and high to low at S/G2M.

### S/G2M fluctuating pluripotency regulators peak in S phase

Next, we set out to determine the S/G2M dynamics of known pluripotency factors. We used curated PluriNet database of 292 pluripotency network components (Som et al., 2010) and searched for their entry in our dataset. Altogether, 97 pluripotency factors were found in our list, enabling us to determine cell cycle profile for 33% of the PluriNet (**Figure 2A, Supplemental Table 2**). Although we did not detect the core pluripotency factors (eg OCT4, SOX2 or NANOG), probably due to no nuclear enrichment (Hurrell et al., 2019), we were able to measure the abundances of several critical pluripotency regulators including SALL4, ESRRB, FBXO15, REST, SALL1, ZFP57, DPPA4 and STAT3, and establish their cell cycle profile in mESCs. In addition to TFs, many of the detected proteins are part of a larger pluripotency network (van den Berg et al., 2010; Gagliardi et al., 2013), and belong to chromatin-modifying complexes largely involved in mediating transcriptional repression (**Supplemental Table 2**). Correlation clustering (k-means = 3) distributed 21 pluripotency regulators into Cluster 1 (S-peak), 45 into Cluster 2 (constant) and 31 into Cluster 3 (G2M-peak) (**Figure 2B**). Pluripotency factors highlighted (bold) in the table below (**Figure 2B**) were among the 3^rd^ Q in Figure 1C, and were considered fluctuating (**Figure S2A**).

**Figure 2.**
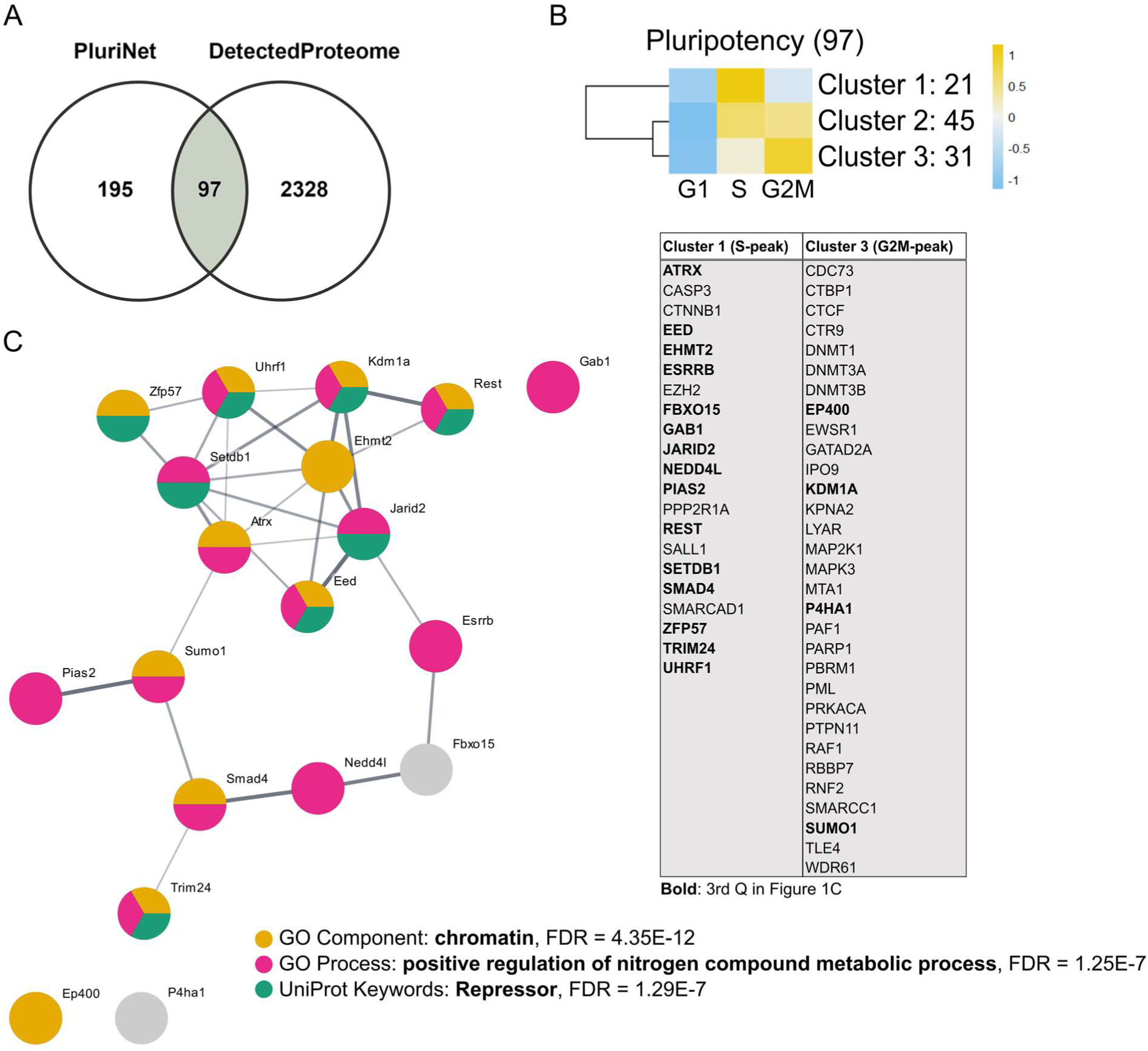
S/G2M fluctuating pluripotency regulators peak in S phase. **A**. Venn diagram showing the overlap between pluripotency network (PluriNet, Som et al., 2010) and the proteins quantified in this study. **B**. Clustered expression heatmap showing the distribution of 97 pluripotency regulators into three major clusters: S-peaking (Cluster 1), Constant (Cluster 2) and G2M-peaking (Cluster 3). Proteins in Clusters 1 and 3 are shown below. Proteins in the 3rd Q in Figure 1C are marked in bold. Note predominant peaking of fluctuating proteins in S phase (15 in S versus 4 in G2M). **C**. Protein interaction network of fluctuating pluripotency regulators derived from Cytoscape/stringApp analysis using edge-weighted layout. Three proteins (EP400, P4HA1, GAB1) were not connected to the network. Overrepresented GO terms were derived via STRING enrichment analysis. See also **Figure S2** and **Supplemental Table 2**.

Several TFs critical for mESC pluripotency including ESRRB, REST, ATRX and ZFP57 as well as components of the Polycomb repressor complex (eg EED, EHMT2, JARID2) and E3 protein ligases (eg NEDD4L, PIAS2, UHFR1) were fluctuating between S and G2M, and their abundance almost exclusively peaked in S phase (15 out of 19) (**Figure S2B**). Pluripotency regulators not changing abundance between S and G2M (constant) included TFs (RBPJ, CTBP2, HELLS) and subunits of the NuRD transcriptional repressor complex (CHD4, HDAC2, MTA2) (**Figure S2C**). Moderately changing TFs included DPPA4, SALL1, SALL4, STAT3, CTNNB1 (all S-peaking), and moderately changing proteins mediating DNA methylation included DNMT1, DNMT3A, DNMT3B (G2M-peaking) and DNMT3L, TET1 (S-peaking) (**Figure S2D**).

STRING network and enrichment analysis of the fluctuating pluripotency factors revealed overrepresentation of GO terms ‘chromatin’ and ‘positive regulation of nitrogen compound metabolic process’, and strong association between proteins assigned with ‘repressor’ activity (**Figure 2C, Supplemental Table 2**). These results reveal variable cell cycle expression profile for different groups of TFs and protein complexes in mESCs and suggest that S-phase peaking of some key pluripotency TFs and Polycomb repressors could reflect fine coordination between cell cycle dynamics and pluripotent identity.

### Inhibition of CDK1 activity induces accumulation of pluripotency regulators in G2M

Given the intimate connection between pluripotency state and cell cycle dynamics, and direct modulation of pluripotency by CDK1 in mESCs aside from its classical role in cell cycle regulation (Neganova et al., 2014; Wang et al., 2017), we set out to determine the sensitivity of pluripotency regulators to perturbation of CDK1 activity. mESC cell cycle fractions treated with CDK1 inhibitor (CDK1i, 10 uM RO3306, 2 hours) were each measured in a 1:1 sample mix using a common standard (SM, Setup 2, **Figure 1A, see Experimental Procedures, Supplemental Table 3**).

On a global scale, similar ratio of proteins had relatively lower abundance (negative z-score) in G1 both in untreated and CDK1i-treated cells (ratio 0.98 vs 0.83, **Figure S3A and S3B**) confirming little impact of CDK1 in G1. Also, the ratio of proteins with positive z-scores was similarly reduced in S and G2M upon CDK1i-treatment (from 0.87 to 0.66 in S, and from 0.88 to 0.73 in G2M), suggesting regulation of global abundances by CDK1 in S/G2M (**Figure S3A and S3B**). Importantly, mitotic proteins showed unchanged cell cycle dynamics upon CDK1i treatment (**Figure S3C**), confirming that CDK1 inhibition did not induce general aberrations in mESC cell cycle. Analysis of the response of 596 globally fluctuating proteins to CDK1i showed that roughly half of the S- and G2M-peaking proteins shifted peak between S and G2M (**Figure S3D**). CDK1i-sensitive proteins in S-peaking set were related to ‘transcription’, and in G2M-peaking set to ‘oxidation-reduction’ and ‘metabolic process’ as revealed using GO analysis (**Figure S3E**), while the CDK1i-insensitive processes were related to DNA damage and repair in S phase, and cell division in G2M (**Figure S3E**), suggesting critical role of CDK1 in balancing functionally relevant processes between S and G2M in mESCs in addition to classical cell cycle regulation.

Next, to determine the balance of pluripotency factor expression during cell cycle in response to CDK1i treatment, we compared proteins via grouping into constant, moderately changing and fluctuating as in Figure S2A-D (**Figure 3A**). This analysis revealed general reduction in the ratio of proteins with positive z-scores in S phase upon CDK1 inhibition. Interestingly, similar pattern was seen for ‘constant’ and ‘moderate’ groups of proteins in G2M but not for the ‘fluctuating’ group where the ratio of proteins with positive z-score increased from 0.26 to 0.74 upon CDK1i treatment (blue rectangle in **Figure 3A**), suggesting pluripotency factor accumulation in G2M when CDK1 activity was inhibited. Indeed, a considerable fraction of S-peaking pluripotency regulators (0.44) showed peak shift from S to G2M along with inhibition of CDK1 (**Figure 3B**), resulting in the reorganisation of S/G2M expression profile (**Figure 3C and 3D**).

**Figure 3.**
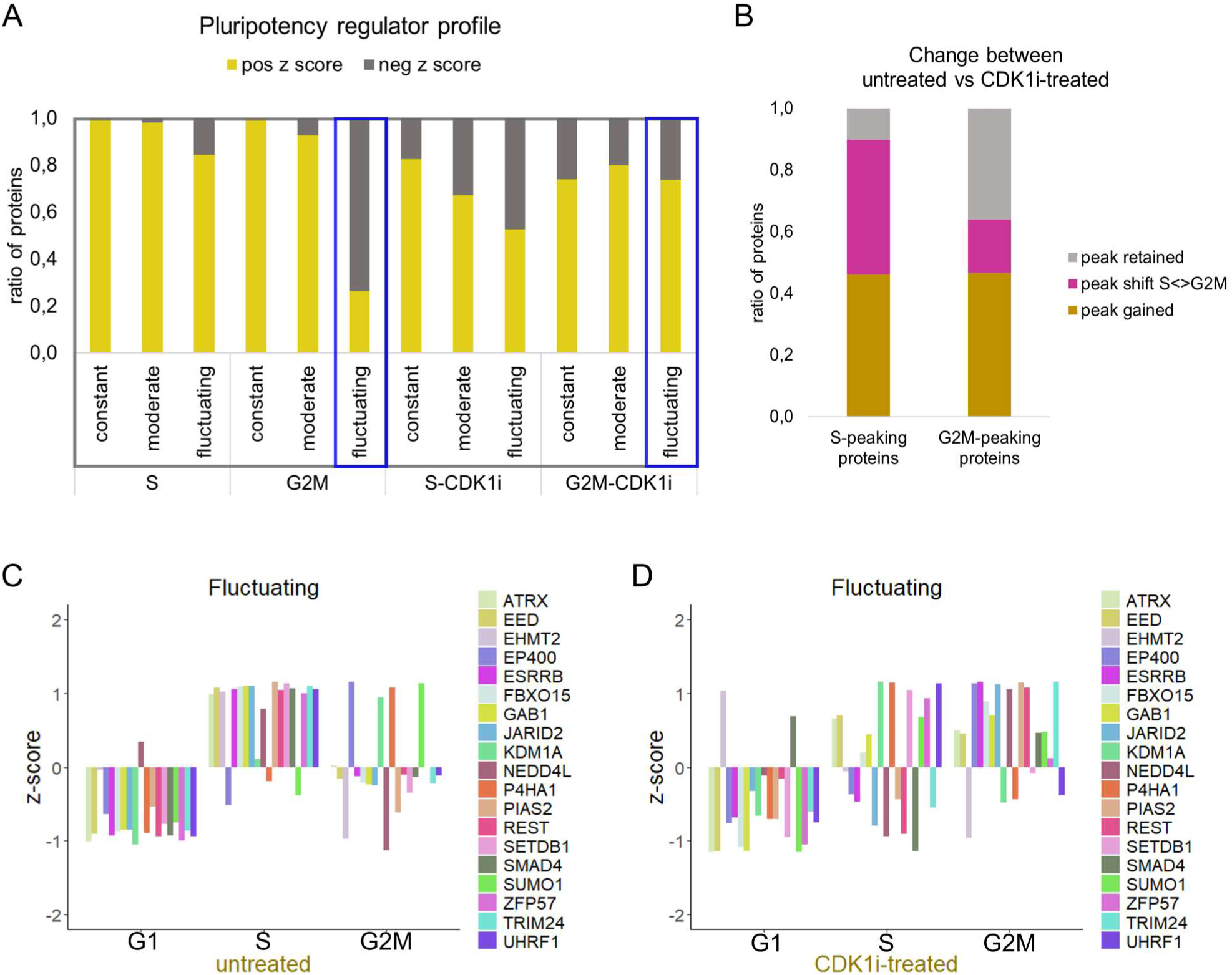
CDK1i induces accumulation of S/G2M fluctuating pluripotency regulators in G2M. **A**. Ratios of pluripotency regulators with either positive or negative z score across untreated or CDK1i-treated cell cycle fractions. Note the increase in the fraction of fluctuating pluripotency regulators with positive z score in G2M phase upon CDK1i-treatment (blue rectangle). **B**. S- and G2M-peaking proteins (from Figure S3B) grouped by change between untreated and CDK1i-treated mESCs. **C and D**. Barplots showing expression dynamics of the fluctuating pluripotency regulators in untreated (C) and CDK1i-treated (D) mESCs between cell cycle phases. Note the reorganisation of expression profiles in S and G2M. See also **Figure S3** and **Supplemental Table 3**.

### Differential sensitivity to inhibition of CDK1

Lastly, we visualised three groups of fluctuating pluripotency factors according to cell cycle profiles to enable a more detailed investigation of changes between S and G2M (**Figure 4A-C**). TFs ESRRB and REST, chromatin regulators JARID2 and TRIM24, and E3 ligases NEDD4L and PIAS2 all switched their maximal levels from S to G2M upon CDK1i (**Figure 4A**). Another group of regulators including ATRX, EED, FBXO15 and GAB1 lost the characteristic peak in S and gained uniform expression between S and G2M when CDK1 activity was inhibited (**Figure 4B**). A few pluripotency regulators including SETDB1, ZFP57 and UHFR1 were insensitive to CDK1i and retained high S to G2M ratio (**Figure 4C**).

**Figure 4.**
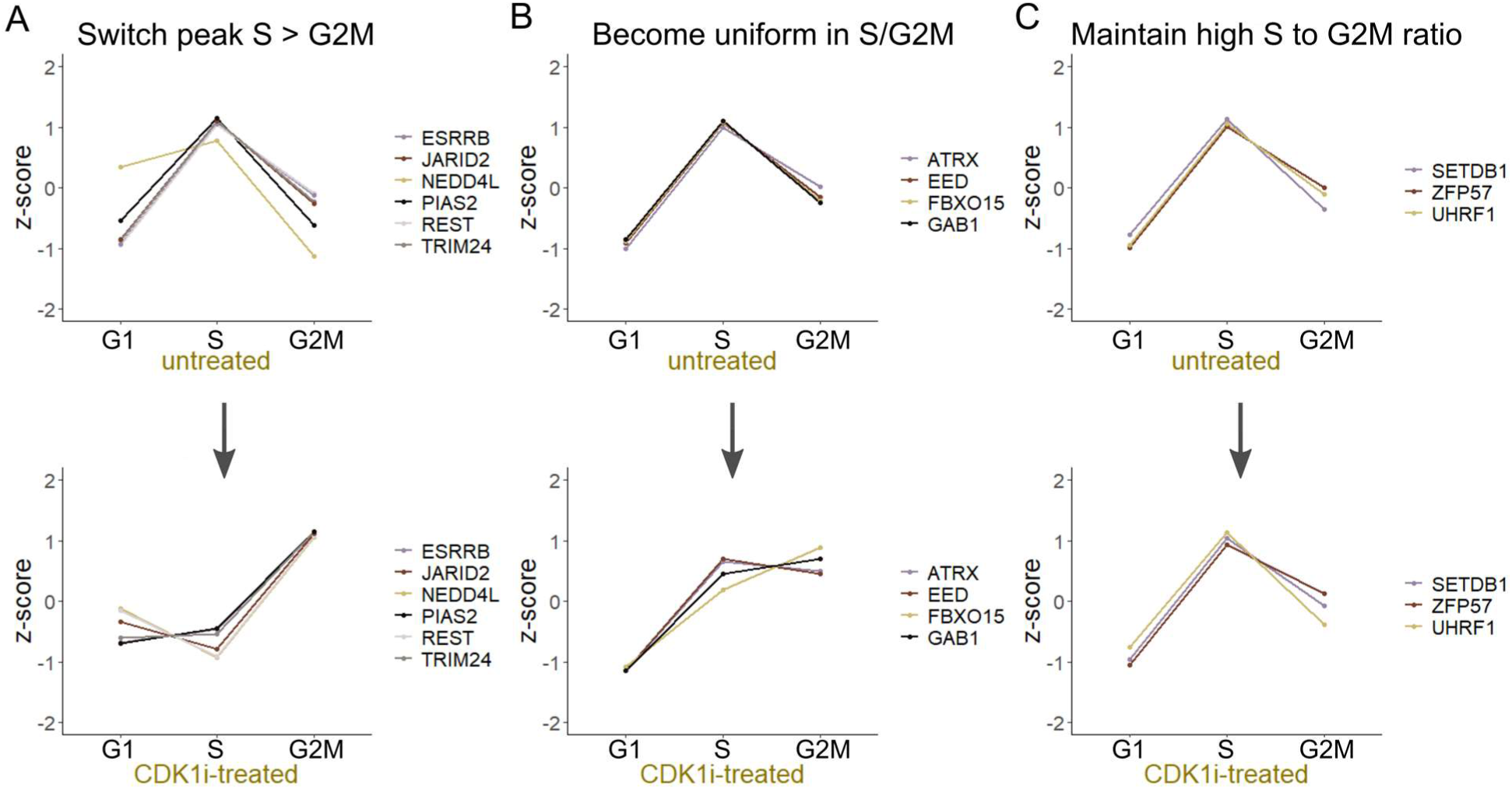
Differential sensitivity of pluripotency regulators to CDK1i treatment. **A-C**. Lineplots showing expression dynamics of different groups of pluripotency regulators either switching peak from S to G2M (eg ESRRB, JARID2, REST in A), becoming uniformly expressed in S and G2M (eg EED, FBXO15 in B) or being insensitive to CDK1 inhibition (eg SETDB1, ZFP57 in C). See also **Supplemental Table 3**.

Together, our results reveal that a large fraction of pluripotency factors change expression between S and G2M, and lose a characteristic S-peaking in the case when CDK1 activity is inhibited.

## DISCUSSION

### S/G2M dynamics of pluripotency regulators

Cell cycle phasing determines alternative ESC pluripotency states and fate choices (Gonzales et al., 2015; Jääger et al., 2019; Van Oudenhove et al., 2016; Pauklin and Vallier, 2013). In this study, we show that mESC proteome is finely balanced between S and G2M phases, and that this balance becomes disrupted upon loss of CDK1 activity.

Chromatin regulators along with tightly regulated TF circuits play important roles in balancing self-renewal and pluripotency in ESCs (Orkin and Hochedlinger, 2011). We found the majority of S/G2M fluctuating pluripotency factors to peak in S phase. These included both critical TFs and transcriptional co-regulators. Interestingly, many factors with repressive activity were found to be particularly sensitive to cell cycle changes. The repressor proteins may be critical to deposit repressive marks temporarily after DNA replication in S phase to prevent differentiation (Petruk et al., 2017), after which their levels need to be reduced again to enable plasticity. The same proteins that we identified here in whole cell lysates were recently found to be over-represented in the chromatin-associated fraction of the mESC proteome (van Mierlo et al., 2019). It is probable then, that via nuclear enrichment, many more pluripotency regulators including core TFs could be profiled using similar approach.

### Reorganisation of S/G2M balance upon loss of CDK1 activity

CDK1 is essential for mESC self-renewal (Zhang et al., 2011) and downregulation of CDK1 causes pluripotent stem cell accumulation in G2 phase, loss of pluripotency and induction of differentiation (Neganova et al., 2014). Therefore, our finding that key pluripotency TFs and co-regulators shift their maximal expression from S to G2M when CDK1 activity is inhibited, suggest S-peaking to have an essential role in maintaining the pluripotent identity of mESC. Not surprisingly then, mESCs spend most of their time in S phase (Stead et al., 2002). Indeed, we found key pluripotency TF ESRRB to switch peak expression from S to G2M upon loss of CDK1 activity. The reduction in expression of ESRRB marks mESC commitment to differentiation (Festuccia et al., 2018). Moreover, variable ESRRB levels separate distinct pluripotency states in G2M (Jääger et al., 2019). Therefore, loss of S-phase peaking of ESRRB protein upon CDK1 inhibition may suggest that the maintenance of naive pluripotency or commitment to differentiation is decided in S phase.

Finally, we propose that downregulation of pluripotency markers observed in response to CDK1 inhibition in bulk ESCs (Neganova et al., 2014) may reflect the reorganisation of abundance profiles between S and G2M observed in this study. Future investigations should address the molecular details of CDK1-mediated control of protein expression at late cell cycle phases and its role in maintaining and/or releasing pluripotent identity of ESCs.

## Author Contributions and Notes

S.K. and K.J. designed research; S.K., R.S., M.S. and K.J. performed research; K.J. and S.K. analyzed data; K.J. wrote the paper with contributions from S.K. and M.L.

This article contains supplemental information online: Supplemental_Information.pdf

## Acknowledgments

We thank Eve Toomsoo and Reet Kurg for assistance with cell sorting, and Mihkel Örd and Viljar Jaks for helpful discussions. We also thank Ian Chambers for kindly sharing the N7-mESC cell line. This research was supported by the Mobilitas Pluss Programme grant (MOBTP28) from the European Regional Development Fund. M.L. and R.S. were supported by ERC Consolidator Grant nr. 649124 PHOSPHOPROCESSORS (M.L.), Centre of Excellence for “Molecular Cell Technologies” TK143 (M.L.), Estonian Science Agency grants IUT2-21 and PRG550 (M.L.). The authors declare no conflict of interest.

