## Supplemental_Information for "Cell cycle-balanced expression of pluripotency regulators via cyclin-dependent kinase 1"

**Supplemental Figures**

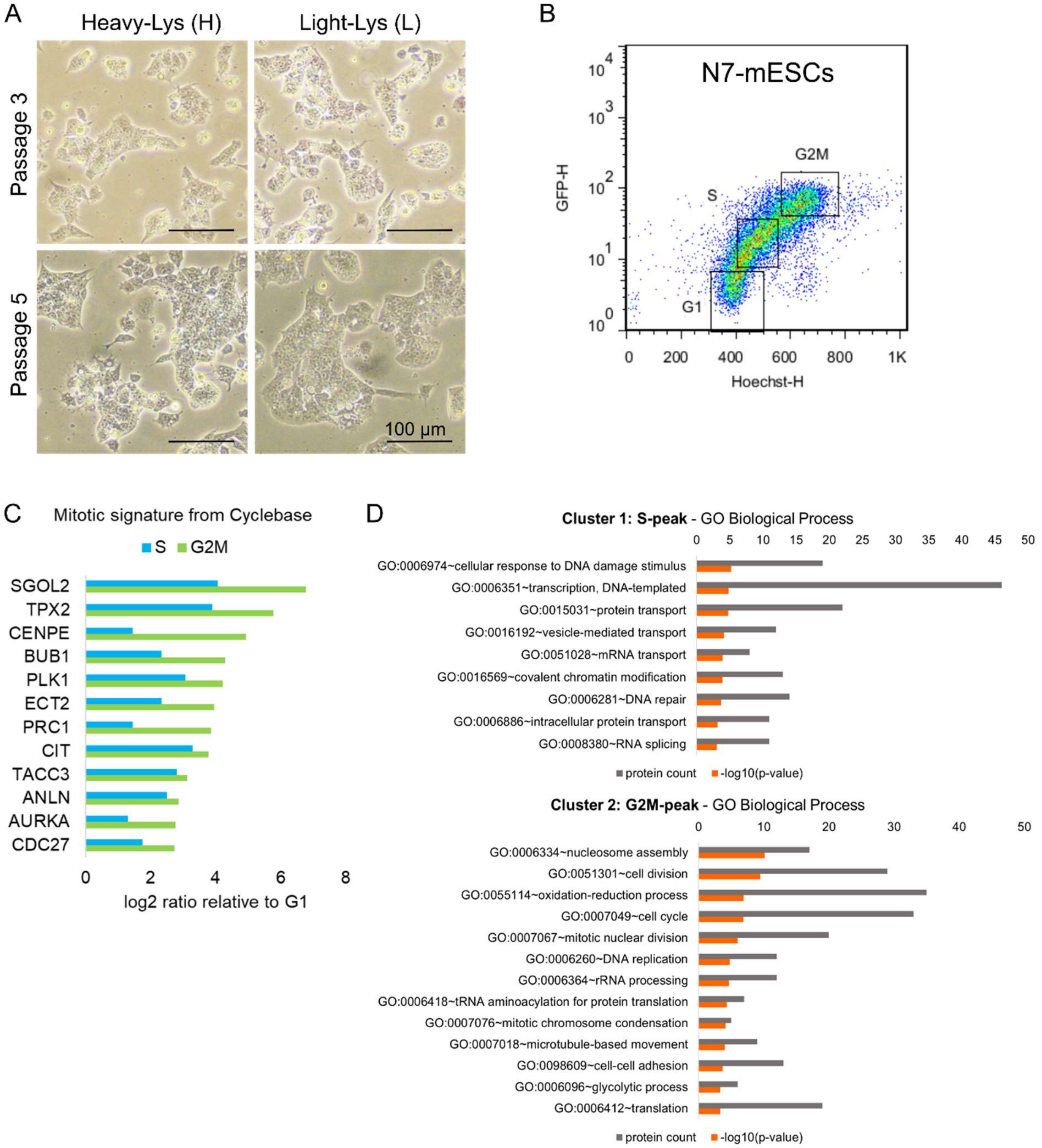


**Figure S1. Cell cycle sorting and characterisation of sorted mESCs. Related to Figure 1 and Experimental Procedures.**

**A.** Brightfield microscope images of N7-mESCs cultured in SILAC media after 3 and 5 passages. **B.** Cell cycle sorting of N7-mESCs using flow cytometry based on Hoechst DNA staining and Cyclin-B1:GFP intensity. **C.** Abundances of mitotic signature proteins in S and G2M phases relative to G1. **D.** Gene ontology Biological Process terms enriched in S-peaking and G2M-peaking sets in mESCs by clustering in Figure 1D.

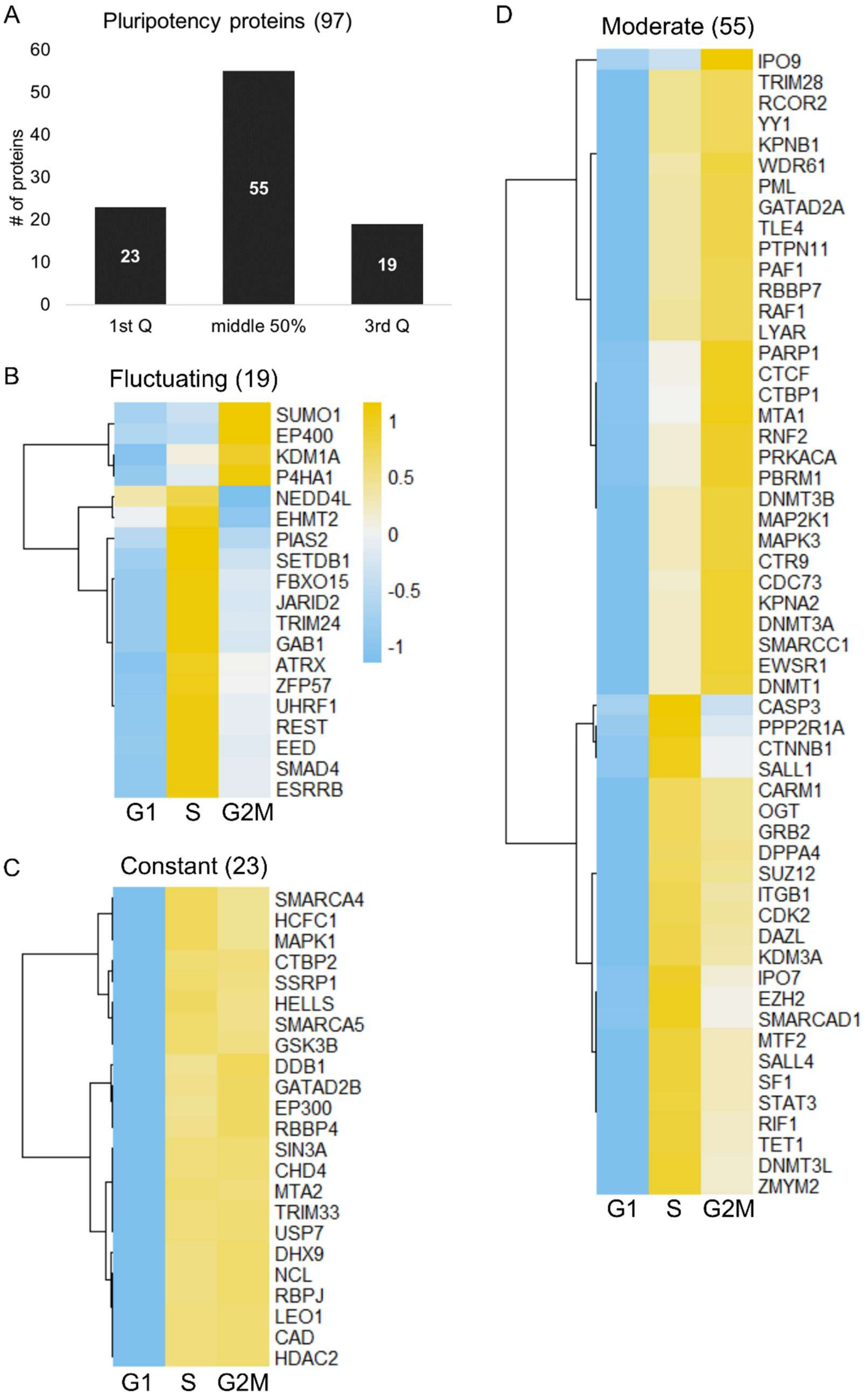


**Figure S2. Cell cycle profiling of pluripotency factors. Related to Figure 2.**

**A.** Distribution of identified pluripotency regulators between boxplot quartiles in Figure 1C. **B-D.** Clustered heatmaps showing scaled abundances of 19 fluctuating (B), 26 constantly expressed (C) and 55 moderately changing (D) pluripotency regulators in mESCs between S and G2M.

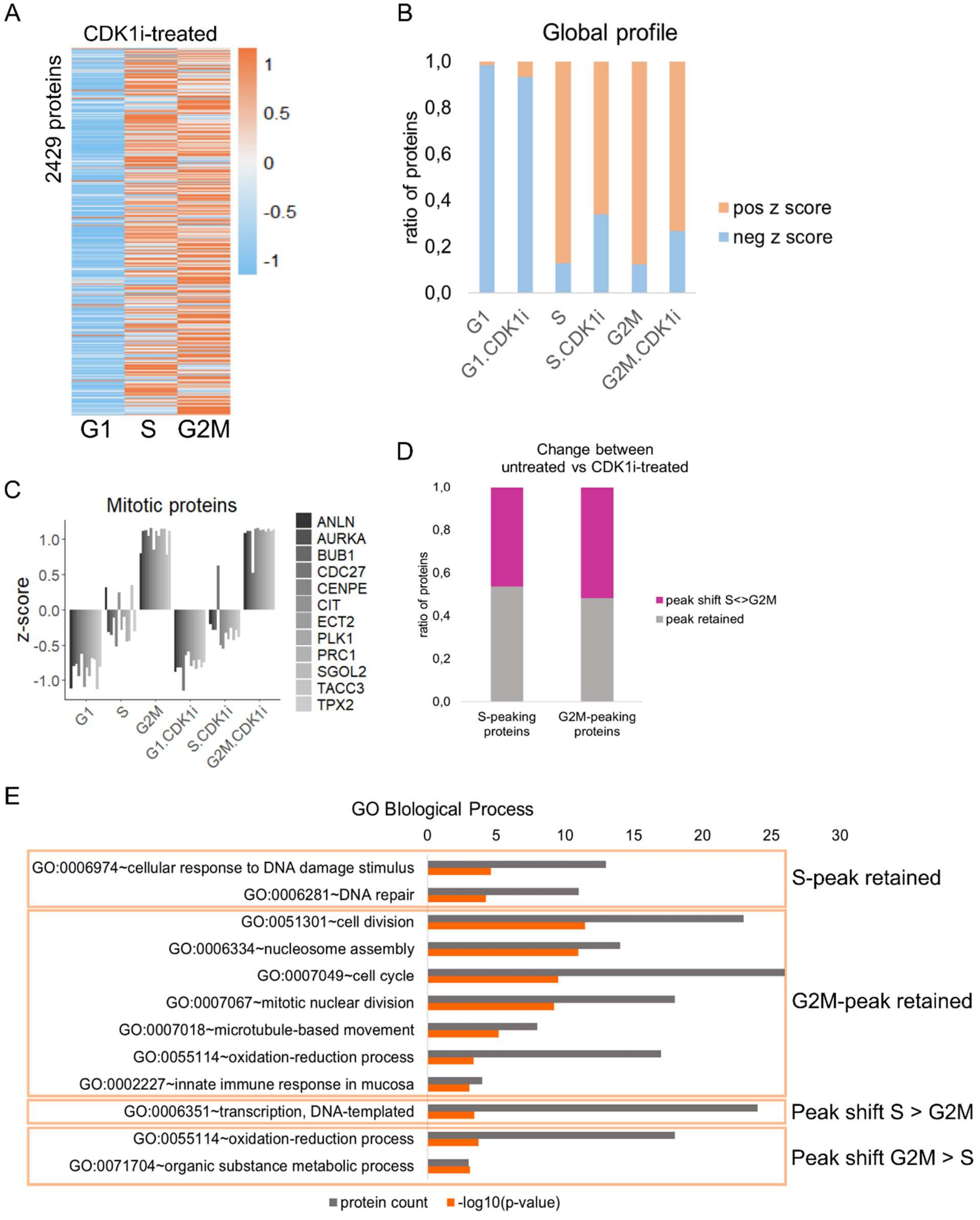


**Figure S3. Cell cycle profile of CDKi-treated mESC proteome. Related to Figure 3.**

**A.** An unclustered heatmap showing the scaled global abundances in CDK1i-treated mESCs (RO3306 10uM, 2 hours) across cell cycle phases. **B.** Ratios of proteins with either positive or negative z score across untreated or CDK1i-treated cell cycle fractions. **C.** Relative abundance profiles of mitotic proteins across cell cycle phases in untreated and CDK1i-treated mESCs. **D.** S- and G2M-peaking proteins (from Figure 1D) grouped by differences between untreated and CDK1i-treated mESCs. **E.** Gene ontology terms enriched in S-peaking and G2M-peaking sets by grouping in D.

### Supplemental Experimental Procedures

#### Proteomics sample preparation

Cells were suspended in 10 volumes of 4% SDS, 100 mM Tris-HCl pH 7.5, 100 mM DTT lysis buffer. Samples were heated at 95ºC for 5 min, followed by probe sonication (Bandelin) (60x 1 s pulses, 50% intensity). Unlysed material were cleared with centrifugation at 14 000 g for 10 min, although, lysis was noted to be nearly complete. 30 µg of protein was precipitated with 2:1:3 (v/v/v) methanol:chloroform:water. Protein pellets were suspended in 25 µl of 7 M urea / 2 M thiourea 100 mM ABC solution, followed by disulfide reduction and cysteine alkylation with 5 mM DTT and 10 mM chloroacetamide for 30 min each at room temperature. Proteins were predigested with 1:50 (enzyme to protein) Lys-C (Wako Chemicals) for 4 h, diluted 5 times with 100 mM ABC and further digested with trypsin (Sigma Aldrich) overnight at room temperature. Peptides were desalted with in-house made C18 StageTips.

#### Nano-LC/MS/MS measurement

Samples were injected to an Ultimate 3000 RSLCnano system (Dionex) using a C18 trap-column (Dionex) and an in-house packed (3 µm C18 particles, Dr Maisch) analytical 50 cm x 75 µm emitter-column (New Objective). Peptides were eluted at 200 nl/min with a 5-35% B 240 min gradient (buffer B: 80% acetonitrile + 0.1% formic acid, buffer A: 0.1% formic acid) to a Q Exactive Plus (Thermo Fisher Scientific) mass spectrometer (MS) using a nano-electrospray source (spray voltage of 2.5 kV). The MS was operated with a top-10 data-dependent acquisition strategy. Briefly, one 350-1400 m/z MS scan at a resolution setting of R=70 000 at 200 m/z was followed by higher-energy collisional dissociation fragmentation (normalized collision energy of 26) of 10 most intense ions (z: +2 to +6) at R=17 500. MS and MS/MS ion target values were 3e6 and 5e4 with 50 ms injection times. Dynamic exclusion was limited to 70 s.

#### Raw Data Processing

Mass spectrometric raw files were processed with MaxQuant software package (version 1.5.6.5). Labelling state (multiplicity) was set to 2 and Lys8 was defined as the heavy amino acid. Methionine oxidation, glutamine/asparagine deamidation and protein N-terminal acetylation were set as variable modifications, while cysteine carbamidomethylation was defined as a fixed modification. Search was performed against UniProt (www.uniprot.org) Mus musculus reference proteome database using the tryptic digestion rule (including cleavages after proline). Only identifications with minimally 1 peptide 7 amino acids long were accepted and transfer of identifications between runs was enabled. Protein quantification criteria were set to 1 peptide with minimally 2 MS1 scans per peptide. Peptide-spectrum match and protein false discovery rate (FDR) was kept below 1% using a target-decoy approach. All other parameters were default.

#### Profiling, enrichment analysis and visualisation of proteomics data

The quantitative information from dual-SILAC measurements was integrated via either common S phase in setup 1 or via common super-mix (SM) in setup 2 to enable comparison between phases. Protein intensities were normalised to S phase replicate 1 (Setup 1) or SM replicate 1 (Setup 2). Only proteins that were identified in all five standard samples (two S phase replicates and three SM replicates) were retained (2434 proteins). For plotting S phase intensities, we used the average of two replicates.

To determine highly fluctuating proteins between S and G2M, the S/G1 and G2M/G1 ratios were calculated, log transformed and standard deviation (stdev) computed. 5 proteins missing both in G1 and G2M were additionally removed, leaving 2429 proteins for expression analyses. Proteins that formed the 3rd (upper) quartile on box plots statistics of stdev-s were defined as highly fluctuating between S and G2M. The 1st (lower) quartile were defined constant, and all others were defined as moderately changing. These proteins were profiled in a similar manner in CDK1i-treated samples using S-CDK1i/G1-CDK1i and G2M-CDK1i/G1-CDK1i ratios.

For profiling of known mitotic marker proteins, we used Cyclebase signature based on both 'peaktime' and 'phenotype' associated with M phase (Santos et al., 2015). For protein interaction network analyses, we used STRING database (Snel et al., 2000; Szklarczyk et al., 2019) via Cytoscape/stringApp (Doncheva et al., 2019). Network layouts were created with 'edge-weighted spring embedded' algorithm applied to all nodes according to STRING score, and nodes colored by GO terms derived via STRING enrichment in Cytoscape (Shannon et al., 2003). Boxplots were generated with BoxPlotR web-tool (http://shiny.chemgrid.org/boxplotr/). Heatmap visualisation and k-means correlation clustering were performed in R using *pheatmap* (https://cran.r-project.org/web/packages/pheatmap/index.html). Venn diagram was generated using Venny 2.1.0 (https://bioinfogp.cnb.csic.es/tools/venny/). Barplots and lineplots were constructed either in Excel or R using *ggplot2*.
